## Supplementary material for "Environmental and molecular noise buffering by the cyanobacterial clock in individual cells": All supplementary info

### Supplemental Figures

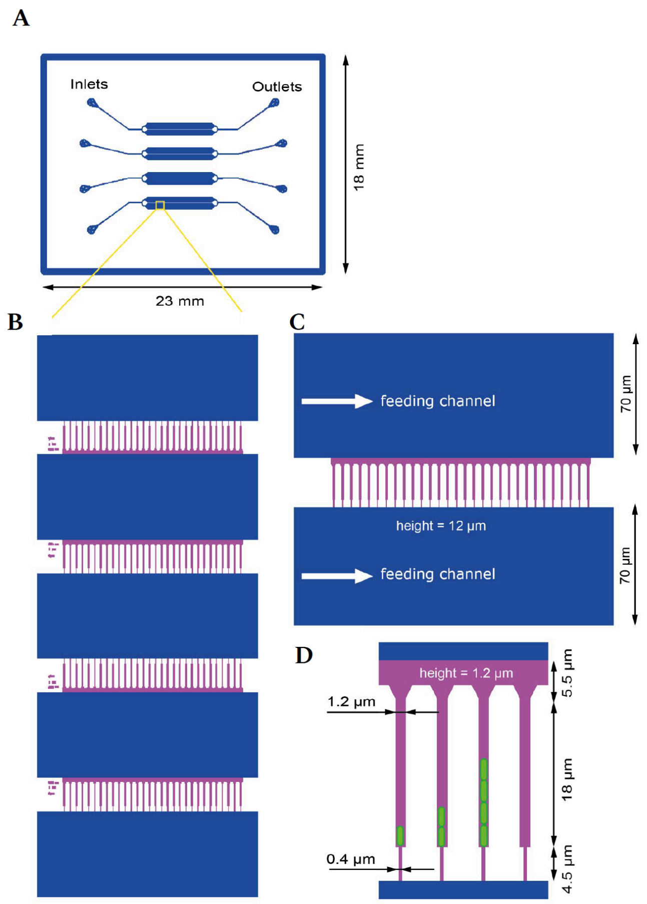


***Figure S1.1: GMM chip design. A:*** *A schematic diagram of the GMM with the sizes of characteristic features indicated. The chip is described in Sachs et al.*[1] ***B-D****: A magnified view of the key chip features. Cells grow from top to bottom of the growth channel and eventually get washed away by the media flow in the feeding channel (indicated by white arrows).* ***D:*** *A magnified view of the growth channels. The key advancement of this MM design comes from the thin channels connecting the growth channels to the top of the neighbouring feeding channel. These connections aid cell loading and facilitate media exchange and waste product removal from the growth channels.*

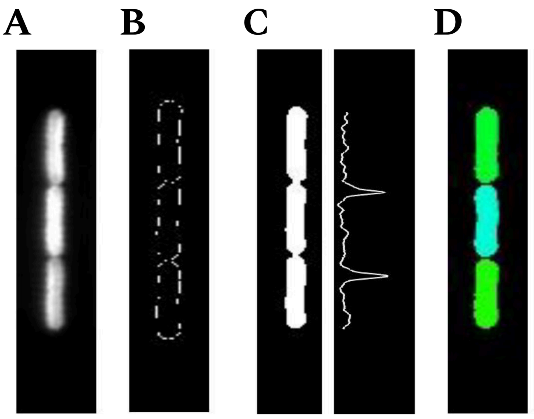


***Figure S1.2: GMM image analysis.*** *Key steps in the image segmentation pipeline: an autofluorescent image (acquired using an RFP filter) (**A**) is used to detect cell outlines (**B**), which are then transformed into an image mask (**C**); the pixel intensity profile along the y-axis is used to separate individual cells from each other (**D**). The code was adapted from Schwall et al.*[2].

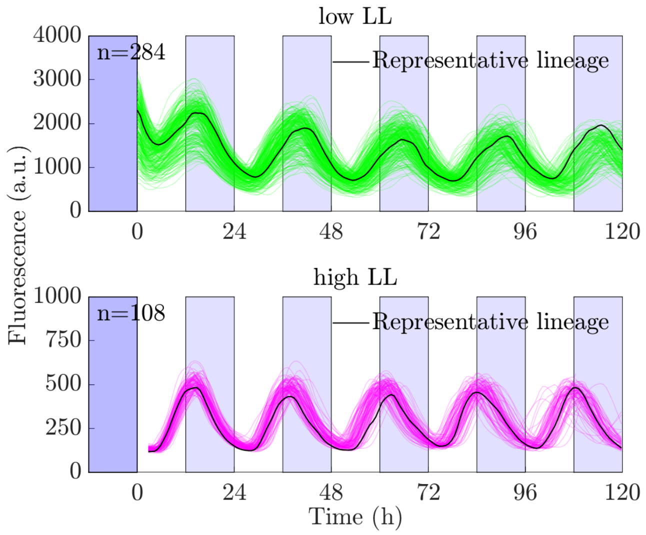


***Figure S1.3 Clock reporter fluorescence under low and high LL regimes**. Individual reporter traces are shown (green and magenta), and a representative trace (black) is highlighted. n indicates the number of cell lineages analysed. Low LL conditions (top panel) correspond to 10 µmol $m^{-2}$ $s^{-1}$ and high LL (bottom panel) correspond to 40 µmol $m^{-2}$ $s^{-1}$. Doubling times were 21.6 ± 5.8 $h^{-1}$ for low LL and 5.1 ± 1.7 $h^{-1}$ for high LL. The differences in absolute fluorescence levels are caused by the differences in the respective imaging set-ups (see **Tab.S1** for details).*

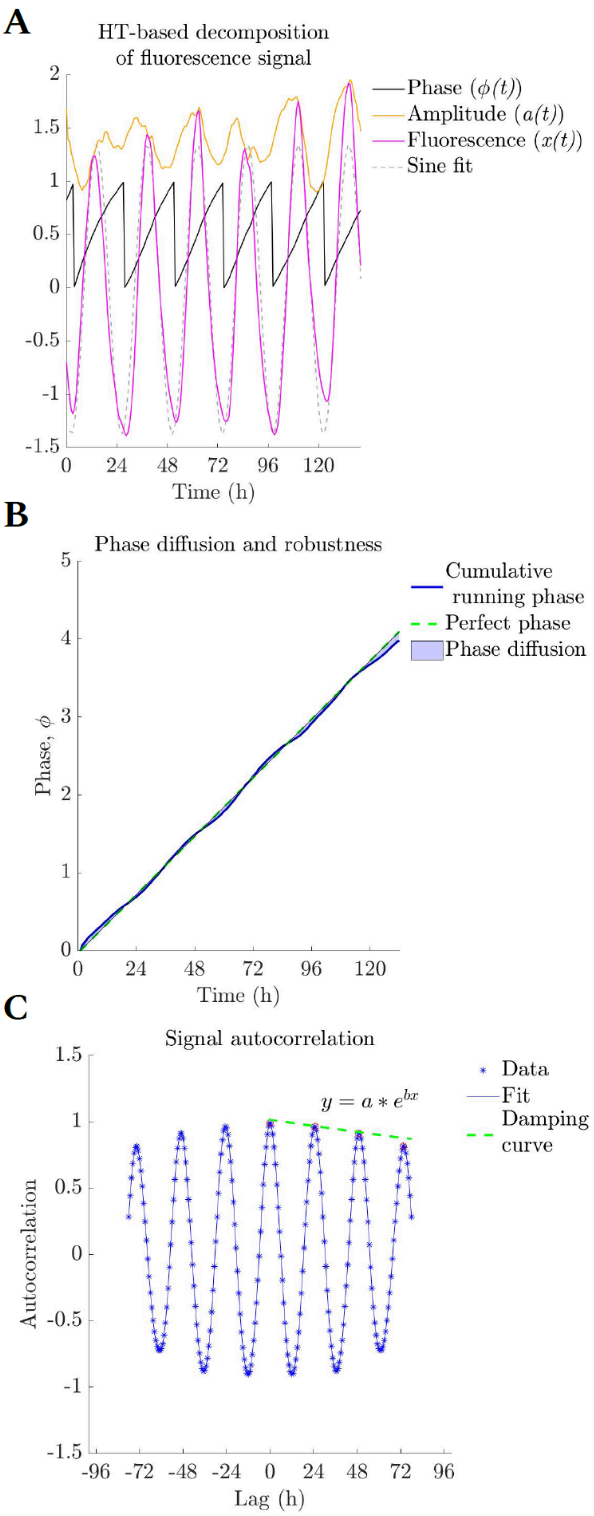


***Figure S1.4 Method to quantify the robustness of clocks within cell lineages. A:*** *Oscillatory clock output was analysed by subtracting the mean of each trace and normalising by its standard deviation (magenta line). Hilbert transform was then used to compute the analytical signal and decompose the clock output into an instantaneous phase (black) and amplitude (yellow). Oscillations were close to sinusoidal waves (dashed blue line).* ***B:*** *The instantaneous phase, running from 0 to 1 every clock period, is added to create a cumulative running phase (blue line). The deviation between the cumulative running phase and a 'perfect' phase (green dashed line) is driven by phase diffusion (light blue shade).* ***C:*** *Signal*

*autocorrelation was used to quantify periods, damping and correlation times of individual lineages. See **Methods** for calculation details.*

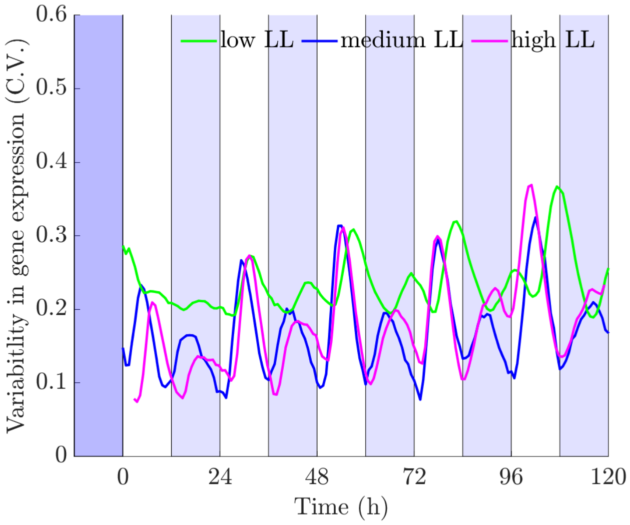


***Figure S1.5 Clock gene expression noise is circadian under LL.** The variability in gene expression (C.V.) had two peaks and one trough per clock cycle. The peaks in C.V. occured at transitions between troughs and peaks in the reporter expression. The pattern was consistent across LL regimes of different intensities. Low LL conditions corresponded to 10 μmol $m^{-2}$ $s^{-1}$, medium LL corresponded to 20 μmol $m^{-2}$ $s^{-1}$, and high LL corresponded to 40 μmol $m^{-2}$ $s^{-1}$.*

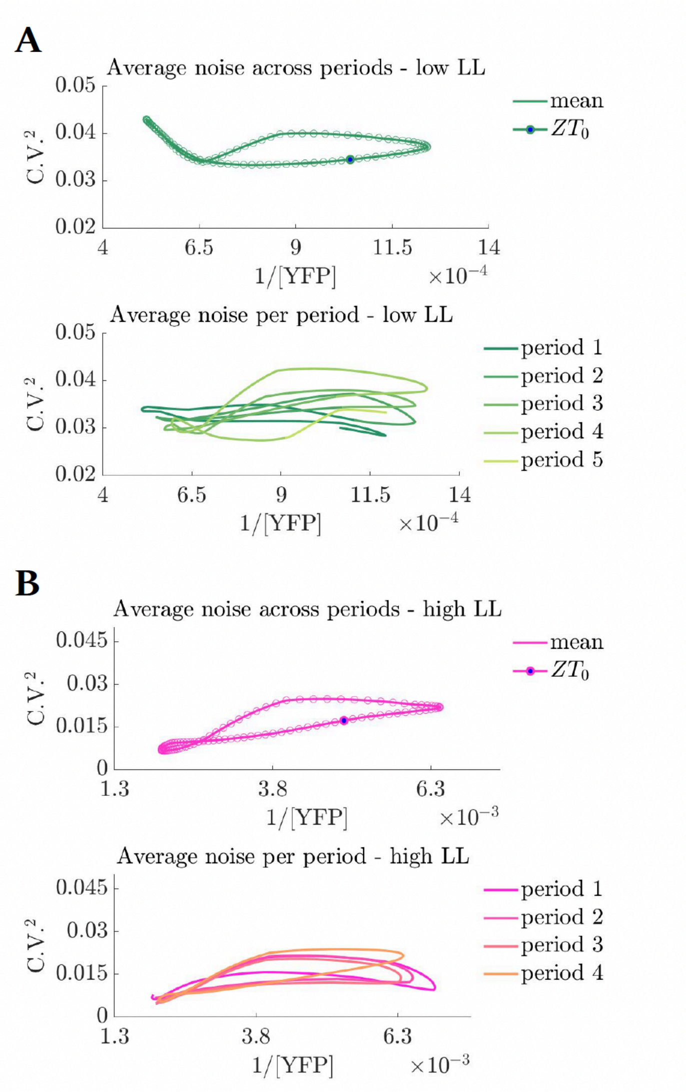


***Figure S1.6 Noise loops in clock amplitude are consistent across clock periods under different LL regimes. A:*** *The top panel represents a noise loop based on the mean reporter oscillation across the experiment and includes intra-lineage variability, whereas the bottom panel has average noise loops per oscillation (period). $ZT_0$= zeitgeber time 0, subjective dawn. Period 1 corresponds to the 1$^{st}$ period after entrainment. Low light data is shown.* ***B:*** *Same as (A), but for high LL. Both panels illustrate that amplitude noise is consistent across periods, and low in all conditions tested. Low LL condition corresponds to 10 μmol $m^{-2}$ $s^{-1}$ and high LL corresponds to 40 μmol $m^{-2}$ $s^{-1}$. To compare the data obtained under different conditions, fluorescence data are background (average fluorescence of an empty channel,* ***Methods****) subtracted.*

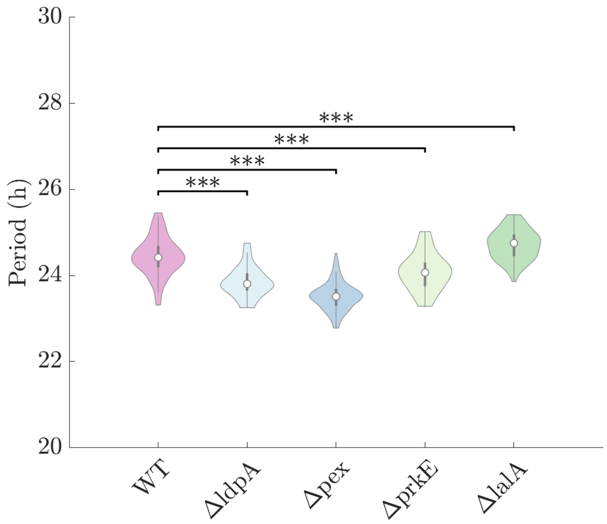


***Figure S2.1 Free running period distributions of KO mutants differ from WT.*** *Cells were observed under medium LL (20 µmol $m^{-2}$ $s^{-1}$). Distribution of periods estimated by computing the autocorrelation function of the fluorescent signal of each lineage (see* ***Methods****). Open circles represent the median and dark bars represent the range between the 1st to 3rd quartiles. Significant differences in the medians of the distributions according to the Wilcoxon rank sum test are represented with an asterisk ( ***, $p<=10^{-3}$).*

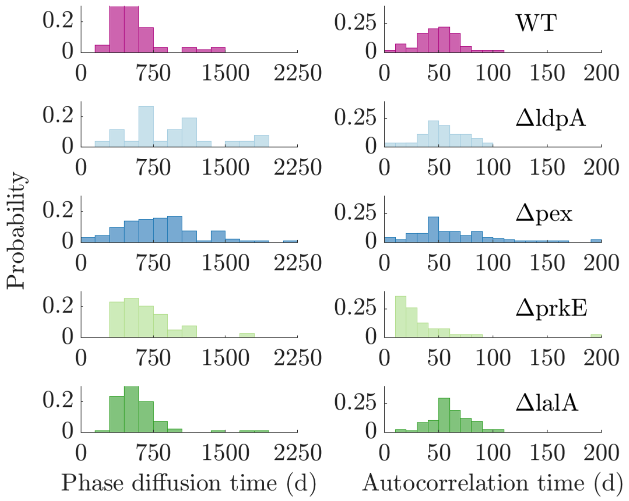


***Figure S2.2 Rhythm stability analysis in regulator KO strains under LL.*** *The clock's phase diffusion time (left column) and autocorrelation time (right column) remain high in the absence of the regulators indicated. The strains were imaged under medium LL (20 µmol $m^{-2}s^{-1}$).*

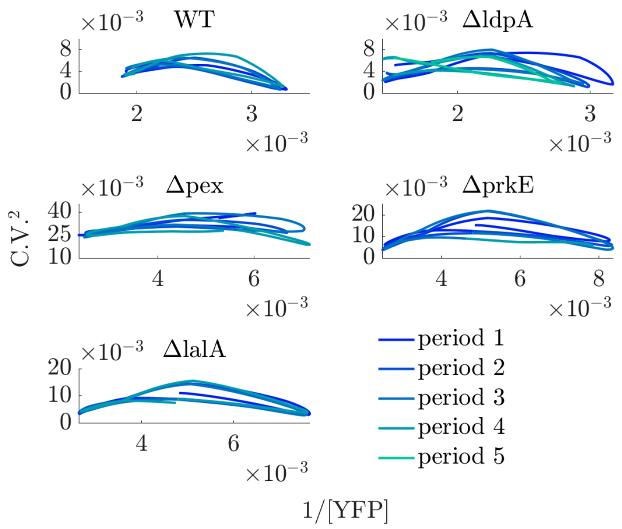


***Figure S2.3 Amplitude noise analysis in regulator KO strains under LL.*** *Noise loops of all the KO mutants largely retain the WT shape, magnitude, and stability across the duration of the experiment. Period 1 corresponds to the 1st period after entrainment. The strains were imaged under medium LL (20 µmol $m^{-2}$ $s^{-1}$).*

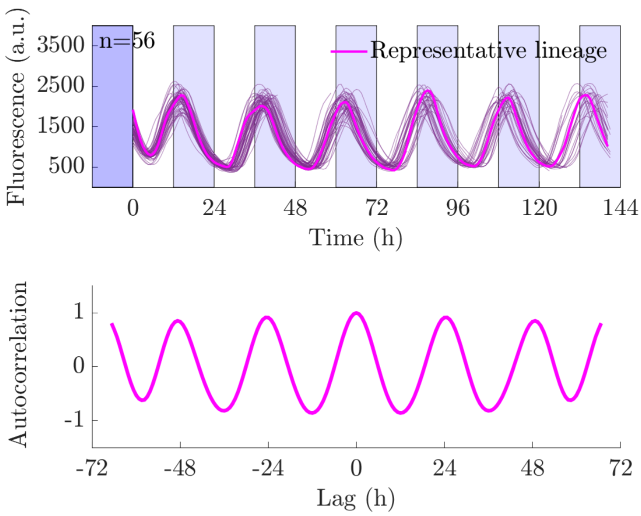


***Figure S3.1 Antibiotic resistance markers were not responsible for the disrupted clock dynamics in KaiC mutants.*** *Clock reporter dynamics are not disrupted in the presence of a gentamicin antibiotic resistance gene (Gent) inserted in neutral site II (NSII) in the WT-Ab strain (see* ***Tab.1****). Individual lineage fluorescence is shown and a representative lineage is highlighted in the top panel. n indicates the number of cell lineages analysed. Bottom panel shows a WT-like autocorrelation.*

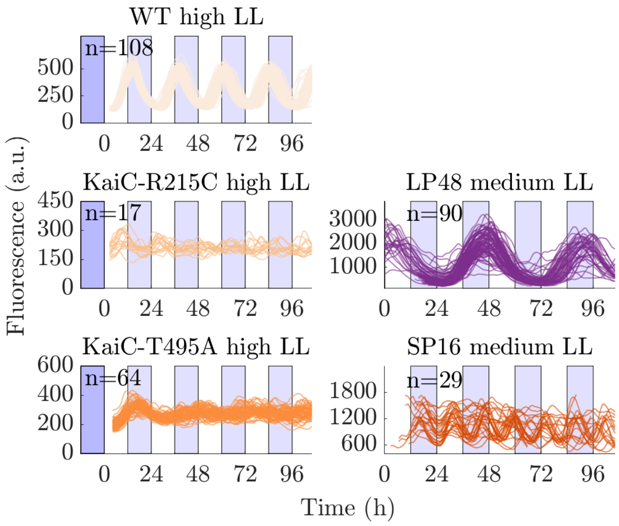


***Figure S3.2 Clock reporter dynamics in KaiC mutant strains under LL.*** *Each line shows an individual cell lineage. n indicates the number of lineages analysed. Medium LL conditions corresponded to 20 μmol m*$^{-2}$ *s*$^{-1}$ *and high LL corresponded to 40 μmol m*$^{-2}$ *s*$^{-1}$*. WT high LL is the same dataset as in Fig.1. SP16 and LP48 were imaged under different conditions. LP48 and SP16 mutants were not entrained, so traces were aligned by the mean time of the first trough for visualisation purposes.*

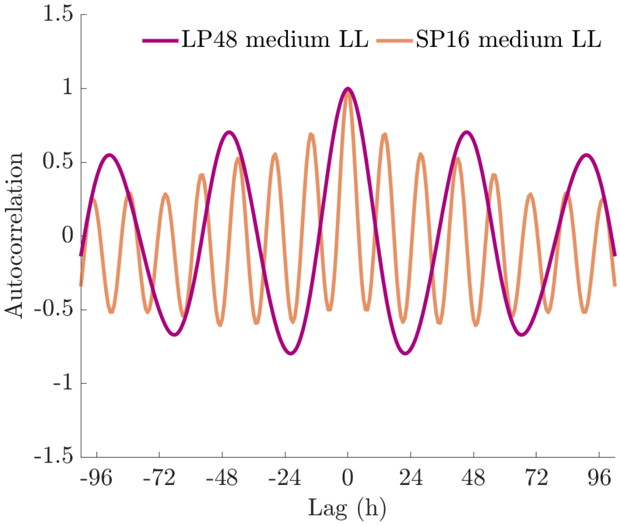


***Figure S3.3 Autocorrelation function in KaiC period mutants.*** *Rhythm autocorrelation of individual lineages shows stronger damping in SP16 than in LP48. SP16, however, goes through more oscillations than LP48 per unit of time. When autocorrelation functions are scaled per period, both clocks show damping at a similar rate (**Fig.3B**).*

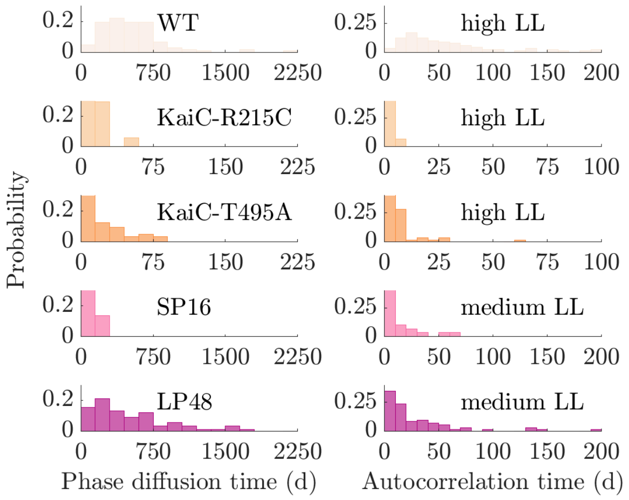


***Figure S3.4 Clock noise is elevated in KaiC mutants.*** *The phase diffusion time (left column) and autocorrelation time (right column) of the free-running clock are shorter in KaiC mutants than in the WT clock. The strains were imaged under medium LL (20 µmol m$^{-2}$s$^{-1}$) and high LL (40 µmol m$^{-2}$s$^{-1}$). Histogram colours as in* ***Fig.3****.*

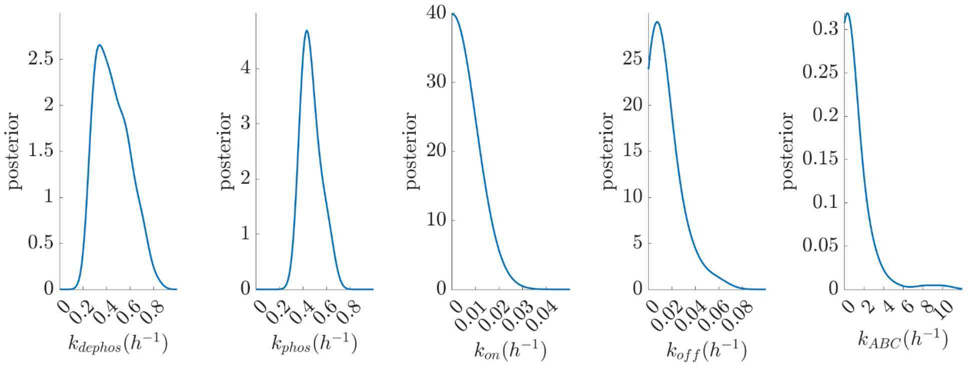


***Figure S3.5 Posterior distributions of model parameters from parameter fitting.*** *ABC-rejection sampling was used with exponential priors spanning several orders of magnitude. 100 best fits were collected and kernel densities were computed. The parameters showed good identifiability except for* $k_{ABC}$.

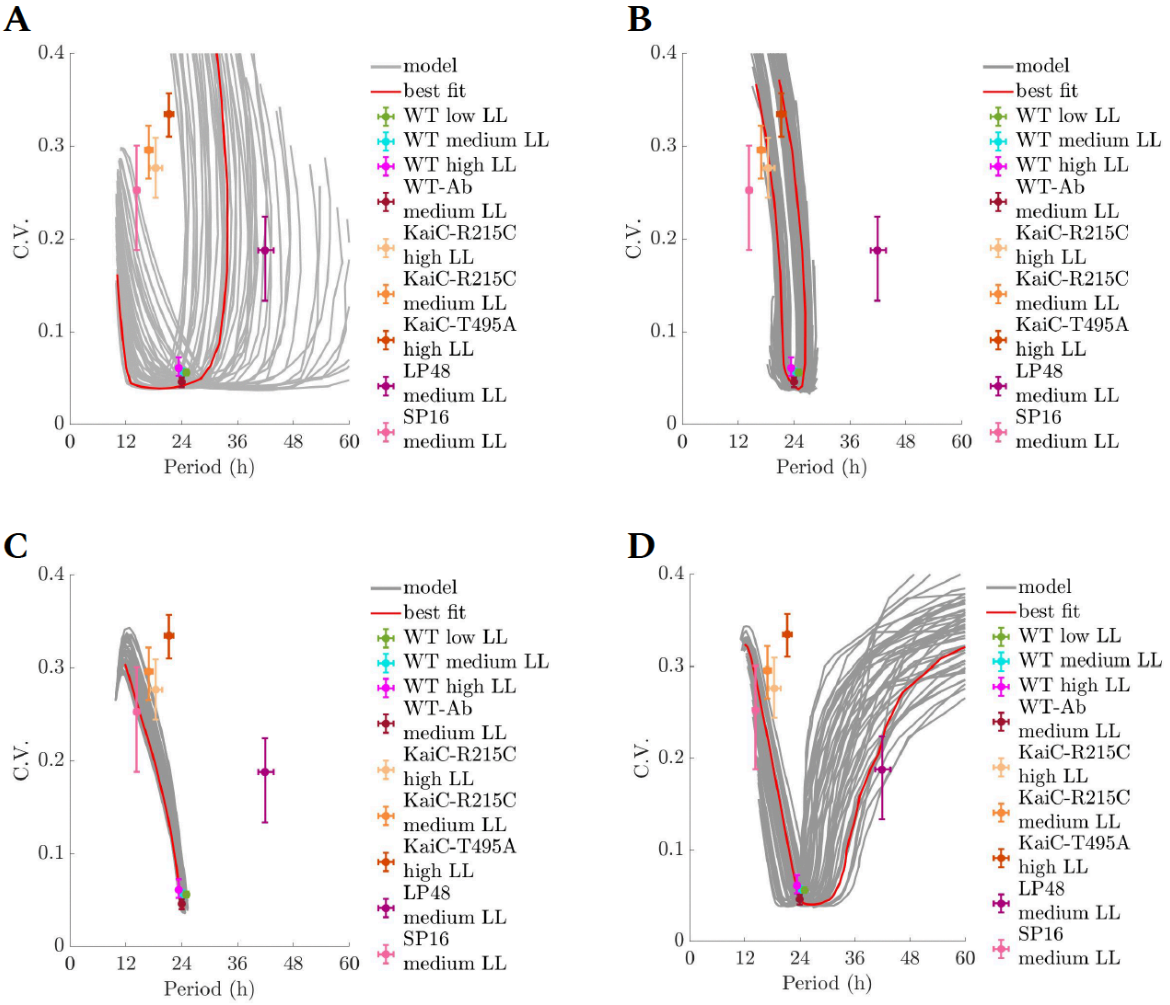


***Figure S3.6 Robustness of the clock to parameter perturbations. A:*** *Perturbations of phosphorylation rate* $k_{phos}$, ***B:*** *KaiA-binding rate* $k_{on}$, ***C:*** *dephosphorylation rate* $k_{dephos}$, *and* ***D:*** *KaiA-unbinding* ($k_{off}$). *Perturbations to the dephosphorylation rate (C) can only speed up the clock. Parameters obtained from ABC-rejection sampling fitted only to WT data in medium LL conditions were varied over several orders of magnitude, displaying a range of mean periods and period noise (obtained from peak times from long lineages simulated for 300 days). The resulting trajectories were smoothed (grey). Perturbation of best fit to WT highlighted as red line.*

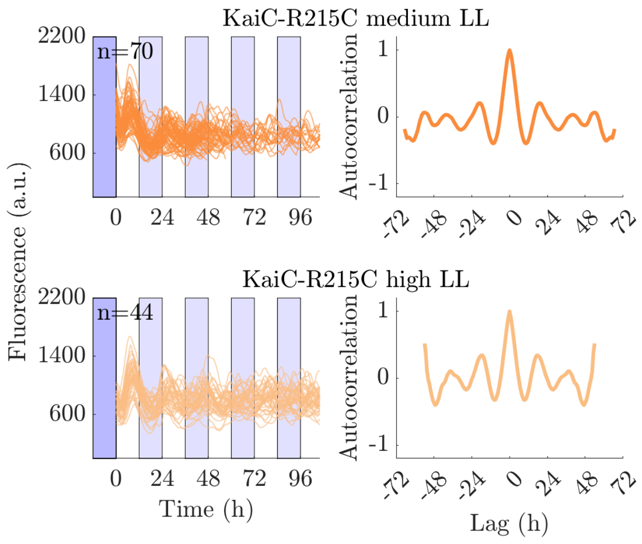


***Fig.S3.7. KaiC-R215C mutant dynamics are maintained across the medium and high light conditions.*** *Autocorrelation of reporter fluorescence is strongly damped, illustrating the elevated noise properties induced by the genetic changes in the mutant.*

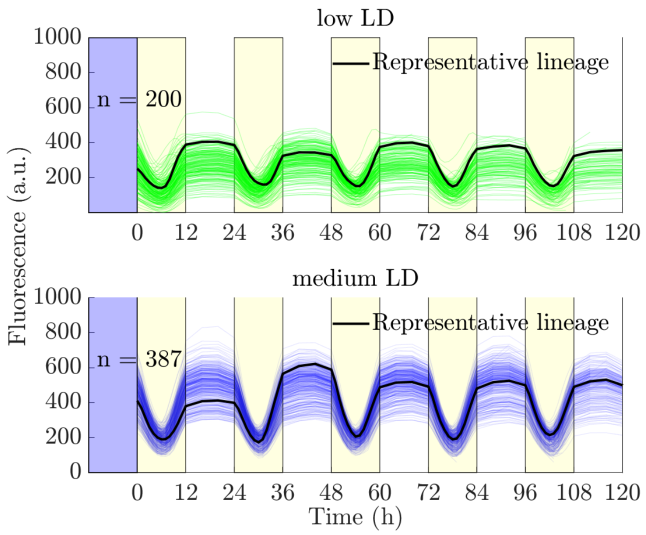


***Figure S4.1 Clock dynamics under 12 h: 12h square LD cycles.*** *Each line shows the fluorescence of an individual lineage, with a representative lineage highlighted in black. n indicates the number of cell lineages analysed. The strains were imaged under low LD (L phase was 10 μmol $m^{-2}s^{-1}$) and medium LL (L phase was 24 μmol $m^{-2}s^{-1}$).*

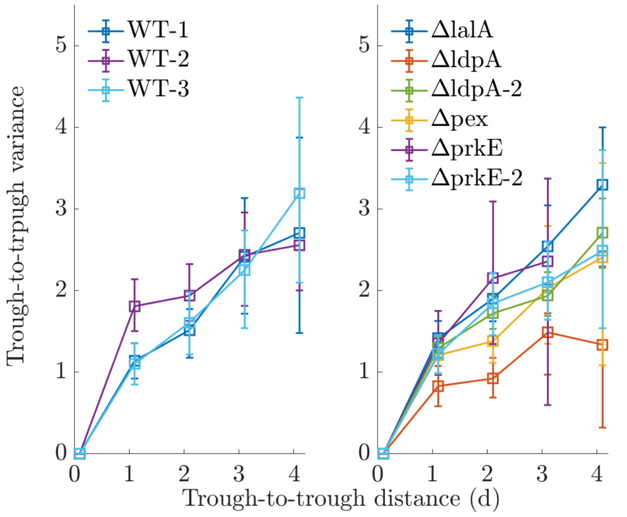


***Figure S4.2 Phase variance under LL in WT (left) and clock regulator-deficient mutants (right).*** *WT repeats (left) demonstrate consistent behaviour of clock phase variance increasing with time. Repeat 1 data were used to train the model and are presented in **Figs.1,3** and **4**. Repeat 2 is shown in Fig.2. Clock regulator data (right) come from Fig.2, apart from ΔldpA-2 (n=211) and ΔprkE-2 (n=42). Error bars in the trough-to-trough variance were obtained through bootstrapping.*

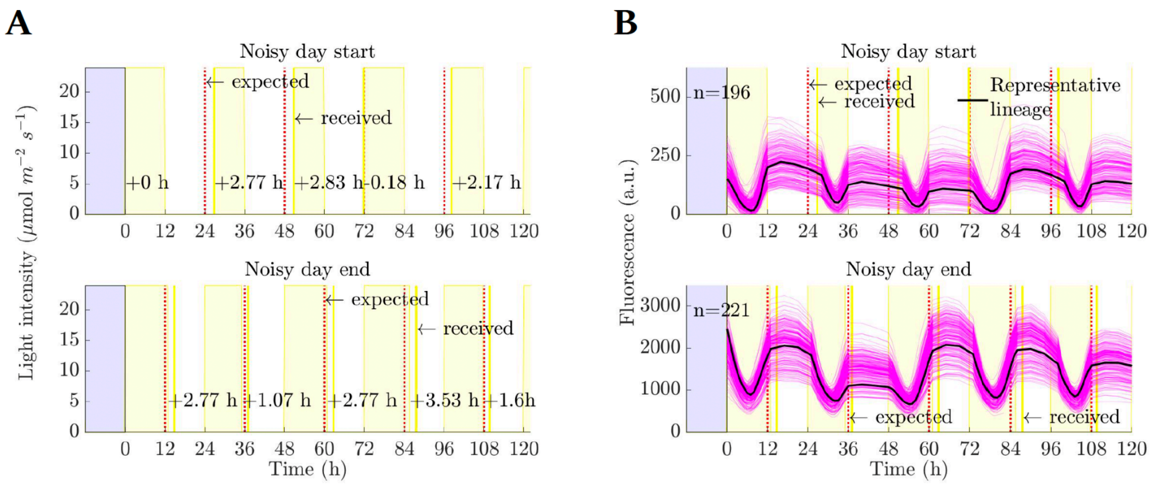


***Figure S5.1 Light conditions and fluorescence reporter data under noisy day start and noisy day end environments. A:*** *Noisy day start (top panel) / end (bottom panel) square LD cycles. The expected L/D switch is shown as a vertical red dashed line, and the real switch is shown as a yellow line. The numbers at the bottom of each graph describe the offset between expected and real L/D switching.* ***B:*** *Clock reporter traces in individual lineages (magenta), with one representative lineage highlighted (black). n indicates the number of cell lineages analysed.*

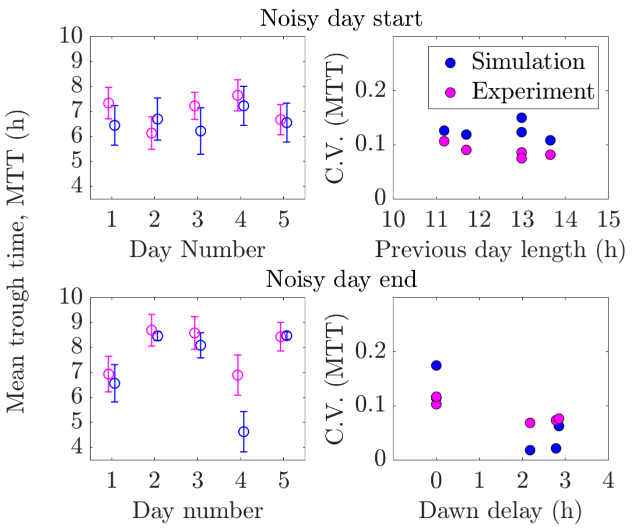


***Figure S5.2 Clock noise is stable across a range of environmental perturbations.*** *Distributions of trough times across cell lineages show comparable noise levels independent of the magnitude of the perturbation (see* ***Fig.5*** *for details). The left column shows the change in trough times across several days. Open circles denote the mean trough times and error bars denote one standard deviation. The right column shows the C.V. of trough time distributions in the two conditions. The noise levels in the experiment are comparable across perturbations but less so in the simulation for 'noisy day end' conditions. Data points in the left column are shifted along the x-axis for visualisation purposes.*

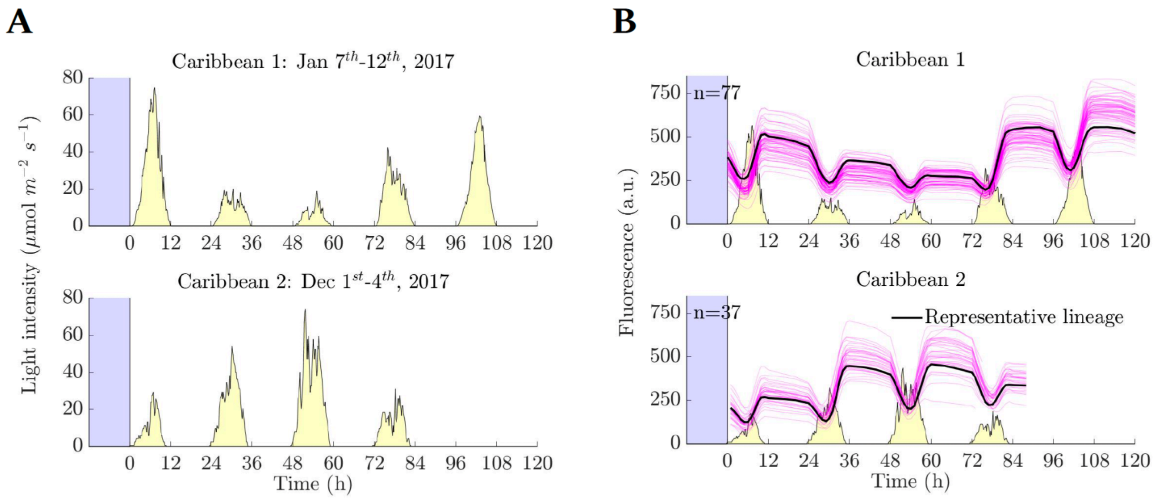


***Figure S6.1 Light conditions (A) and fluorescence reporter data (B) under the Caribbean environments.*** *Clock reporter traces in individual lineages (magenta), with one representative lineage highlighted (black). n indicates the number of cell lineages analysed.*

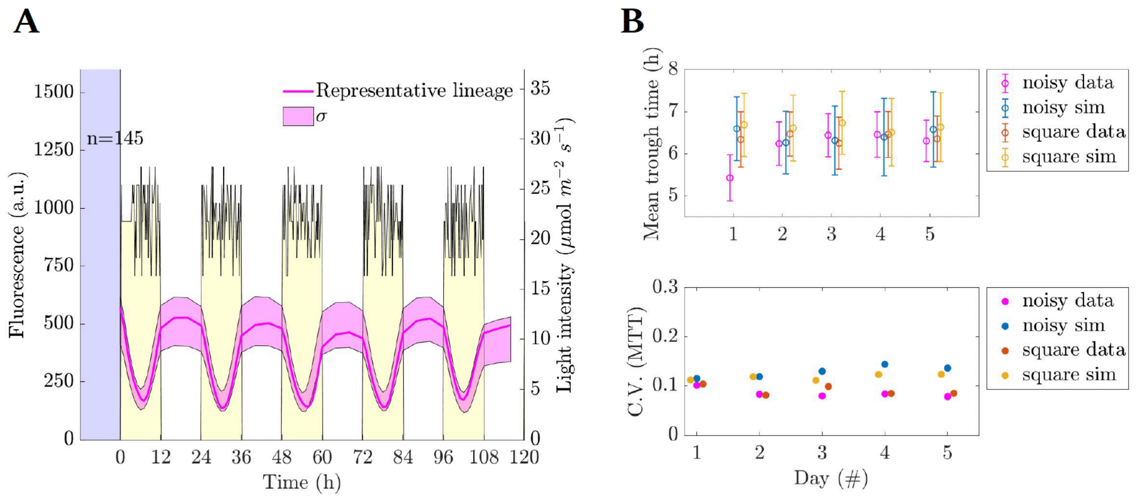


***Figure S6.2. Clock filters high-frequency noise in the environment. A:*** *Cells were exposed to 12 h: 12 h square LD cycles with high-frequency (every 10 min) light fluctuations of up to 25% around the mean light level (24 μmol $m^{-2}s^{-1}$). A representative fluorescence trace and data distribution within one standard deviation (σ) are shown.* ***B:*** *As predicted by the model, the means (top) and C.V.s of mean trough times (MTTs) are similar across days and between noisy and non-noisy LD cycles. Data points for different conditions are shifted along the x-axis for visualisation purposes.*

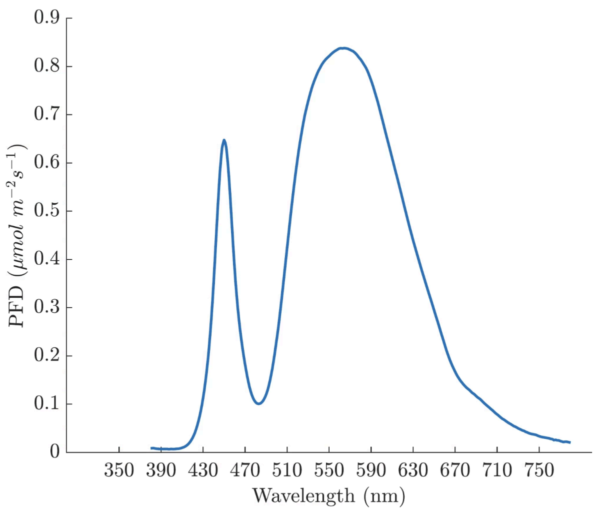


***Figure S7. Light spectrum of circular LED array for ambient illumination in the microscope.*** *The peak of Photon Flux Density (PFD) corresponds to a wavelength λ of 555 nm, and the mean PFD at $\lambda_{380\text{-}780}$ nm was 128.3 μmol $m^{-2}$ $s^{-1}$.*

| Dataset as named in each figure (light condition) | Figure | Camera (CoolSNAP/Prime) | YFP imaging conditions (exposure time, binning) |
|---|---|---|---|
| WT (high LL) | Fig.1,3; Fig.S1.3, S1.5, S1.6, S3.2 | CoolSNAP | 120 ms, 2 |
| WT or WT-2 (medium LL) | Fig.2; Fig.S2.3, S4.2 | CoolSNAP | 120 ms, 2 |
| WT or WT-1 (medium LL) | Fig.1,3,4; Fig.S1.5, S4.2 | CoolSNAP | 100 ms, 2 |
| WT-3 (medium LL) | Fig.S4.2 | Prime | 100 ms, 2 |
| WT (low LL) | Fig.1,4; Fig.S1.3, S1.5; S1.6 | Prime | 100 ms, 2 |
| WT (medium LD) | Fig.4; Fig.S4.1 | Prime | 100 ms, 1 |
| WT (low LD) | Fig.4; Fig.S4.1 | Prime | 100 ms, 1 |
| WT (noisy day start) | Fig.5; Fig.S5.1 | Prime | 100 ms, 1 |
| WT (noisy day end) | Fig.5; Fig.S5.1 | Prime | 100 ms, 2 |
| WT (Caribbean 1) | Fig.6; Fig.S6.1 | Prime | 100 ms, 1 |
| WT (Caribbean 2) | Fig.6; Fig.S6.1 | Prime | 100 ms, 1 |
| WT (noisy day) | Fig.S6.2 | Prime | 100 ms, 1 |
| WT-Ab (medium LL) | Fig.S3.1 | Prime | 100 ms, 2 |
| ΔlalA (medium LL) | Fig.2; Fig.S2.3 | CoolSNAP | 120 ms, 2 |
| ΔldpA (medium LL) | Fig.2; Fig.S2.3 | CoolSNAP | 120 ms, 2 |
| ΔldpA-2 (medium LL) | Fig.S4.2 | Prime | 120 ms, 2 |
| Δpex (medium LL) | Fig.2; Fig.S2.3 | CoolSNAP | 120 ms, 2 |
| ΔprkE (medium LL) | Fig.2; Fig.S2.3 | CoolSNAP | 100 ms, 2 |

| | | | |
|---|---|---|---|
| ΔprkE-2 (medium LL) | Fig.S4.2 | Prime | 100 ms, 2 |
| LP48 (medium LL) | Fig.3; Fig.S3.2 | Prime | 100 ms, 2 |
| SP16 (medium LL) | Fig.3; Fig.S3.2 | Prime | 100 ms, 2 |
| KaiC-R215C (high LL) | Fig.3; Fig.S3.2 | CoolSNAP | 120 ms, 2 |
| KaiC-R215C (medium LL) | Fig.S3.7 | Prime | 100 ms, 2 |
| KaiC-R215C (high LL) | Fig.S3.7 | Prime | 100 ms, 2 |
| KaiC-T495A (high LL) | Fig.3; Fig.S3.2 | CoolSNAP | 120 ms, 2 |

***Table S1. Imaging conditions for all experiments.*** *Only figures reporting reporter fluorescence, noise loops, or additional technical repeats are listed. All YFP images were obtained upon exposure to maximum excitation intensity, as further described in* ***Methods.*** *Binning was performed using an in-built function in Metamorph.*

**Supplemental movies:**

**Movie S1:** A time-lapse movie of a WT strain (from a data set shown in **Fig.2**). The strain expresses a pkaiBC:eYFP:fsLVA transcriptional reporter (shown in green), which displays circadian oscillations. Cells are entrained in the chip upon exposure to 12 h darkness and then observed under free-running, continuous light conditions (medium LL, 20 μmol $m^{-2}$ $s^{-1}$). Phase contrast images are shown in the background in grey. The imaging frequency is every 45 min.

**Movie S2:** A time-lapse movie of a WT strain (from a data set shown in **Fig.4**). The strain expresses a pkaiBC:eYFP:fsLVA transcriptional reporter (shown in green), which displays circadian oscillations. Cells are entrained in the chip upon exposure to 12 h darkness and then observed under square-shaped 12 h: 12 h square LD cycles (medium LD, 24 μmol $m^{-2}$ $s^{-1}$). Phase contrast images are shown in the background in grey. The imaging frequency is every 60 min during the day and every 240 min during the night.

***References:***


1. *Sachs, C. C. et al. Image-Based Single Cell Profiling: High-Throughput Processing of Mother Machine Experiments. PLoS ONE* ***11****, e0163453 (2016).*
2. *Schwall, C. P. et al. Tunable phenotypic variability through an autoregulatory alternative sigma factor circuit. Mol. Syst. Biol.* ***17****, e9832 (2021).*
